# Characterization of p38α signaling networks in cancer cells using quantitative proteomics and phosphoproteomics

**DOI:** 10.1101/2022.09.09.507259

**Authors:** Yuzhen Dan, Nevenka Radic, Marina Gay, Adrià Fernández-Torras, Gianluca Arauz, Marta Vilaseca, Patrick Aloy, Begoña Canovas, Angel R. Nebreda

## Abstract

p38α (encoded by *MAPK14*) is a protein kinase that regulates cellular responses to almost all types of environmental and intracellular stresses. Upon activation, p38α phosphorylates many substrates both in the cytoplasm and nucleus, allowing this pathway to regulate a wide variety of cellular processes. While the role of p38α in the stress response has been widely investigated, its implication in cell homeostasis is less understood. To investigate the signaling networks regulated by p38α in normally proliferating cancer cells, we performed quantitative proteomic and phosphoproteomic analyses in breast cancer cells in which this pathway had been either genetically targeted or chemically inhibited. Our study identified with high confidence 35 proteins and 82 phosphoproteins (114 phosphosites) that are modulated by p38α, and highlighted the implication of various protein kinases, including MK2 and mTOR, in the p38α-regulated signaling networks. Moreover, functional analyses revealed an important contribution of p38α to the regulation of cell adhesion, DNA replication and RNA metabolism. Indeed, we provide experimental evidence supporting that p38α negatively regulates cell adhesion, and showed that this p38α function is likely mediated by the modulation of the adaptor protein ArgBP2. Collectively, our results illustrate the complexity of the p38α regulated signaling networks, provide valuable information on p38α-dependent phosphorylation events in cancer cells, and document a mechanism by which p38α can regulate cell adhesion.

## Introduction

p38α is a ubiquitously expressed protein kinase that belongs to the mitogen-activated protein kinase (MAPK) family and plays a key role in maintaining cellular homeostasis. Essentially all stressful stimuli, including UV light, heat and osmotic shock, nutrient starvation and oncogene activation, as well as inflammatory factors, lead to p38α activation which then controls numerous cellular processes (1–3). p38α is activated through dual phosphorylation of the Thr-Gly-Tyr motif in its activation loop by the MAP2Ks MKK3 and MKK6, which in turn can be phosphorylated by one of at least 8 different MAP3Ks (4). Upon activation, p38α can potentially phosphorylate many proteins on Ser or Thr residues, which are usually followed by a Pro residue. According to PhosphositePlus (5), 316 phosphosites on 179 distinct proteins have been reported to be positively regulated by p38α in different contexts, but only a fraction of these have been validated as direct p38α targets. The p38α substrates are located throughout the cell and can regulate a wide variety of processes, including transcription and chromatin remodeling, mRNA stability and translation, protein degradation and localization, cell cycle progression, endocytosis, metabolism and cytoskeleton dynamics (2,6). In addition to the stress response, p38α has been shown to modulate inflammation and the immune response, and has therefore been considered as a potential target for diseases such as rheumatoid arthritis, chronic obstructive pulmonary disease or asthma (7). There is also evidence implicating p38α activation in the response of cancer cells to chemotherapeutic treatments, and p38α inhibitors have been shown to potentiate the cytotoxic effect of several chemotherapeutic drugs (8). Moreover, the ability of p38α to regulate processes such as cell migration, proliferation, differentiation and apoptosis suggests that modulation of this pathway could be of therapeutic benefit in a wide group of diseases, such as cancer, autoimmune disorders, neurodegenerative diseases and malfunction of the cardiovascular system (3).

Given the physiological importance of the p38α pathway in health and disease, and its potential as therapeutic target, a comprehensive picture of the substrate repertoire and biochemical mechanisms regulated by p38α signaling is imperative. Unlike other MAPK family members such as ERK1 and ERK2, whose phosphoproteome has been extensively studied by using mass spectrometry-based methods (reviewed by (9,10)), few attempts have been made to identify the full complement of p38α substrates in mammalian cells. Previous reports have investigated phosphorylation changes that are potentially controlled by p38 MAPKs in differentiating myoblasts and in early mouse embryos, as well as in U2OS cells treated with the stress stimuli UV or anisomycin. All these studies were based on the use of pyridyl imidazole inhibitors such as SB203580 (11–14). However, to our knowledge, the phosphoproteome regulated by p38α in non-stressed mammalian cancer cells has not been characterized.

To study the signaling networks orchestrated by p38α in the atypical regulatory context of a cancer cell, we have performed global quantitative proteomic and phosphoproteomic analyses in proliferating breast cancer cells. We targeted p38 MAPK signaling using a p38α genetic knockout (KO) or the chemical inhibitor PH797804 (15), which is more selective for p38α than the widely used compound SB203580 that has well-known off-target effects (16–18). By comparing both datasets, we identified a number of proteins and phosphoproteins that can be regulated by p38α, either directly or through downstream kinases, and identified several cellular processes that are essential for the proper function of cancer cells and in which the p38α-regulated proteome and phosphoproteome are implicated. In particular, we predicted and experimentally validated a role for p38α in the regulation of cancer cell adhesion, which we show implicates the adaptor protein ArgBP2. Taken together, our results reveal the complexity of the p38α-regulated signaling networks in cancer cells, including a high confidence resource of phosphorylation events and signaling networks modulated by p38α, and provide new insights into the regulation of cell adhesion by p38α.

## Experimental Procedures

### Cell culture

A cancer cell line was derived from a mouse epithelial mammary tumor expressing the polyoma middle T (PyMT) antigen together with ubiquitin-CreERT2 (UbCre) and floxed alleles of *Mapk14*, the gene encoding p38α (19). Cells were cultured in Dulbecco’s Modified Eagle Medium (Sigma, D5796) supplemented with 10% fetal bovine serum, 1% penicillin/streptomycin (Labclinics, L0022-100) and 1% L-glutamine (Labclinics, x0550-100), at 37°C in a humidified incubator containing 5% CO_2_. Cells were routinely tested for mycoplasma using Mycoplasma Detection Kit (Lonza, LT07-318) according to the manufacturer’s protocol.

### HEK 293T transfection and lentiviral infection

HEK293T cells were plated in order to reach 80% confluence at the moment of transfection. For transfection in a 100 mm dish, 5 μg of target DNA was mixed with 4.5 μg of δ89 and 500 ng of VSVG packaging plasmids in 450 μl of sterile water. Then, 50 μl of 2.5 M CaCl2 were added dropwise to the mixture. After incubating for 5 min at RT, 500 μl of 2xHBS were added dropwise to the mixture, which was then incubated 20 min at RT and added to the cells. Growth medium was changed 16 h after transfection. Lentiviruses were collected 48 h and 72 h after transfection, mixed 1:1 with fresh medium and added to the desired cells together with 8 μg/ml Polybrene (Sigma, TR-1003).

### Generation of ArgBP2 KO cells

ArgBP2 KO cells were generated in the cancer cell line mentioned above (19) using the CRISPR-Cas9 system. In brief, Cas9 cells were generated by infecting with lentivirus expressing Cas9-Blast (Addgene, #52962) followed by blasticidin selection (5 μg/ml) for 72 h. The following sgRNAs targeting *Sorbs2* were cloned into Lenti_sgRNA_EFS_GFP (LRG, Addgene #65656):

sgRNA_*Sorbs2*_Fwd1: CACCGAGACAAAAGATCACCGACCC,
sgRNA_*Sorbs2*_Rev1: AAACGGGTCGGTGATCTTTTGTCTC,
sgRNA_ *Sorbs2*_Fwd2: CACCGGACACCAGGAAATTTCGAT,
sgRNA_*Sorbs2*_Rev2: AAACATCGAAATTTCCTGGTGTCC.

sgRNA oligos were phosphorylated and annealed in a 10 μl reaction containing 10 μM of forward and reverse sgRNA oligos (100 μM stock), T4 Ligation buffer (NEB, B0202) and T4 PNK (NEB, M0201) using the following thermocycler program: 37°C 30 min, 95°C 5 min, ramp down to 25°C at 5°C/min. For plasmid digestion, 5 μg of LRG sgRNA expression plasmid was digested with FastDigest Esp3I (ThermoFisher, FD0454) and gel purified using QIAquick Gel Extraction Kit (Quiagen, 28706X4). The ligation mixture containing 50 ng of digested LRG, 1 μl of diluted (1:200) oligo duplex, 5 μl of 2x Quick Ligase buffer and 1 μl of Quick ligase (NEB, M2200) in a final volume of 10 μl was incubated for 10 min at RT and then used to transform DH5α competent cells. Ampicillin resistant clones were selected, expanded and the plasmid DNA was extracted (Sigma, PNL350). Lentiviruses were generated in HEK293T cells and then used to infect Cas9 cells. Transduced cells, either single GFP^+^ cells (to obtain clones 1 and 2) or 250 GFP^+^ cells (to obtain the pool) were sorted per well into a 96 well plate (BD FACSAria Fusion Cell Sorter). The two clones and the pool were expanded and the decreased ArgBP2 expression was confirmed by immunoblotting.

### Cell attachment assay

24-well plates were either left empty or coated with 500 μl of collagen I (DMEM, 15 mM HEPES, 0.1 mg/ml BSA, 10 μg/ml Collagen I) overnight at 4°C and dried before use. 30,000 cells were plated into each well of pre-coated plates. After 0.5 h and 1 h incubation, non-adherent cells were removed by washing three times with PBS. Adherent cells were fixed for 20 min with 4% paraformaldehyde in PBS and stained with crystal violet for 20 min. After washing once, plates were dried and images were taken using TE200 NIKON (Olympus DP72).

### Cell detachment assay

For cell detachment assays, cells were plated at confluency in 6-well plates. After 24 h, cells were washed with PBS and incubated with trypsin/EDTA (0.05/0.02%) for 8 min. The wells were then filled with medium and non-adherent cells were removed by washing with PBS. Adherent cells were fixed with 4% paraformaldehyde and stained with crystal violet. After a wash step, plates were dried and scanned. The cell-occupied areas were measured using Fiji.

### 3D spheroid formation assay

The spheroid formation assay was performed as described (20). Lids of 10 cm culture plates were seeded with 20 μl of a 10^6^ cells/ml suspension. To prevent dehydration of the drops, 10 ml PBS was added to the bottom of the plate. Cells were incubated at 37 °C and 5% CO2 for 48 h to allow aggregation. Spheroids were visualized using LEICA DMi8 (Orca Flash LT), and the spheroid areas were measured using Fiji.

### Quantitative real-time PCR

Total RNA was obtained using PureLink RNA mini kit (Ambion, 12183018A) and reversely transcribed with SuperScript IV (Invitrogen, 18090010). The obtained cDNA was used as a template for quantitative PCR reactions with SYBR green (ThermoFisher, 4472942) using QuantStudio™ 6 Flex Real-Time PCR System. Fold induction compared with untreated controls was calculated by the delta-delta CT method. qRT-PCR samples were analyzed in triplicate. *Sorbs2* pre-mRNA primers were designed following the instruction from (21). The following primers were used in this study:

*Sorbs2*_Fwd: ACACCCTAAGCTCCAATAAG, *Sorbs2*_Rev: TTCTTGAAGTTCCCACAAAC, *Pre_Sorbs2_Fwd*: TTTTCCCAATGCTCATTCCA, *Pre_Sorbs2*_Rev: CTTGTCTGTTGGTTCCTGGG, *Gapdh_Fwd*: CTTCACCACCATGGAGGAGGC, *Gapdh_Rev*: GGCATGGACTGTGGTCATGAG.

### Immunoblotting

Cells were lysed in lysis buffer containing 50 mM Tris HCl pH 7.5, 150 mM NaCl, 2 mM EDTA, 1% NP40, 2 mM PMSF, 2 mM mycrocystin, 2 mM sodium orthovanadate, 1 mM DTT and 1x EDTA-free complete protease inhibitor cocktail (Roche, 11873580001), for 30 min on ice and centrifuged at 15000 rpm for 10 min at 4°C. Protein concentration was estimated using the RC DC protein assay kit II (BioRad, 5000122) with BSA as reference. Total proteins (20-40 μg) were resolved on 8%-12% gradient SDS-PAGE gels and electroblotted onto nitrocellulose membranes (Whatman, 10401396). Following blocking in 5% non-fat milk dissolved in TBST (20 mM Trizma base, 150 mM NaCl, 0.1% Tween-20) for 1 h at RT, membranes were incubated with primary antibodies at 4°C overnight. After washing with TBST, membranes were incubated with secondary antibodies for 1 h at RT and visualized using Odyssey Infrared Imaging System (Li-Cor, Biosciences). The primary antibody (1:1000) was diluted in 5% BSA in TBST. The following primary and secondary antibodies were used: Anti-goat HSP27 (Santa Cruz, sc-1049), anti-rabbit phospho-HSP27 (Ser82) (Cell Signaling, #2401), anti-rabbit phospho-MK2 (Thr334) (Cell Signaling, #3007), anti-mouse phospho-p38 (Thr180/Tyr182) (BD Biosciences, 612288), anti-mouse p38α (Santa Cruz, sc81621), anti-mouse ArgBP2 (Santa Cruz, sc-514671), antimouse α-Tubulin (Sigma T9026), Alexa-Fluor anti-mouse 680 (ThermoFisher, A21057), Alexa-Fluor anti-rabbit 680 (ThermoFisher, A21076), IRDye 800CW donkey anti-goat IgG (Li-COR, 926-32214)

### Spheroid immunofluorescence

Spheroids were fixed in 4% paraformaldehyde for 45 min and quenched with 20 mM glycine for 20 min at RT. Then, spheroids were permeabilized in 0.5% Triton X-100 for 30 min and blocked in 1% BSA in PBS for 1 h at RT. Samples were incubated with anti-rabbit ZO-1 primary antibody (1:100) (life technologies, 40-2200) at 4°C overnight in a dark chamber followed by incubation with the Alexa-conjugated anti-rabbit 488 secondary antibody (1:500) (Invitrogen) for 1 h at RT in the dark. Then, spheroids were incubated with DAPI for 15 min and washed with PBS. Mounting media (Dako, s3023) was applied to the slides and spheroids were visualized using Zeiss LSM 780 confocal microscope. The mean intensity was quantified using Fiji.

### SILAC experimental design and statistical rationale

Cells were cultured in SILAC medium DMEM (Silantes) supplemented with heavy, light or medium lysine and arginine until at least 95% of proteins were labeled. Heavy-labeled cells were treated with ethanol for 48 h, washed, split, and harvested 48 h later. Light-labeled cells were treated with 100 nM 4-OHT for 48 h to induce p38α deletion, washed, split, and harvested 48 h later. Medium-labeled cells were treated for 96 h with 2 μM PH797804. Samples were then lysed, digested and enriched for phosphopeptides, and were analyzed by nanoLC-MS/MS at IRB Barcelona’s Mass Spectrometry & Proteomics Facility.

Protein intensities were used for the statistical analysis. Within each SILAC experiment, protein quantitation was normalized by summing the abundance values for each channel over all proteins identified within an experiment. All abundance values were corrected in all channels by a constant factor (x 1E9) per channel so that at the end the total abundance is the same for all channels. Data were first transformed to a log scale to apply a linear model. We filtered the data to retain only proteins with valid quantification values in at least 4 of the samples for the total protein dataset and in at least 3 of the samples for the p-sites dataset. Missing values were imputed with normally distributed random numbers (centered at −1.8 standard deviations units and spread 0.3 standard deviations units with respect non-missing values). To adjust for batch effect, a linear model was used with SILAC batch as fixed effect. Model fitting was accomplished with the lmFit function of the limma package (22) of R statistical software (23). Comparison between groups was done (LH, MH, ML) to find out which species significantly changed. For each comparison, estimated fold changes and p-values were calculated. The batch effect was removed from SILAC intensities. The resulting adjusted intensities were filtered with those proteins significant in at least one comparison, according to the FC and p-value, and z-score normalized. In addition, we retained only proteins that out of the seven replicates analyzed showed valid quantification values in at least four replicates for the proteomic dataset or at least three replicates for the phosphoproteomic dataset.

### Protein digestion

Protein samples were quantified using Pierce™ 660 Protein Assay Kit (plus Ionic Detergent Compatibility Reagent, IDCR; Thermo Scientific). Then, 103.3 μg of each heavy, medium and light labeled protein sample were mixed and digested with trypsin, using the FASP (*Filter-Aided Sample Prep*) approach. The recovered peptide solutions were acidified with formic acid (FA) (1% final concentration). The volume of acidified peptide solution was reduced to 300 μL on a SpeedVac vacuum system and desalted in a C18 tip (P200 Toptip, PolyLC), as per manufacturer’s indications. Samples were dried down and redissolved in 1% formic acid for nanoLC-MS/MS analysis or were subjected to phosphopeptide enrichment (600 ng on column).

### Phosphopeptide enrichment

Phosphopeptide enrichment was carried out with TiO2 magnetic beads (GE Healthcare TiO_2_ Mag Sepharose, Instructions 28-9537-65 AB). Briefly, samples were resuspended in loading buffer (1 M glycolic acid in 80% acetonitrile, 5% TFA) and loaded onto the magnetic beads. The beads were washed once with loading buffer and twice with 80% ACN 1% TFA. The phosphopeptides bound to the TiO2 magnetic beads were eluted using ammonium water (pH12). Phosphopeptide enriched fractions were dried down and redissolved in 1% formic acid for nanoLC-MS/MS analysis (600 ng on column).

### NanoLC-MS/MS analysis

Peptides were analyzed using an Orbitrap Fusion Lumos™ Tribrid mass spectrometer (Thermo Scientific) equipped with a Thermo Scientific Dionex Ultimate 3000 ultrahigh-pressure chromatographic system (Thermo Scientific) and an Advion TriVersa NanoMate (Advion Inc. Biosciences) as the nanospray interface. C18 trap (300 μm i.d x 5 mm, C18 PepMap100, 5 μm, 100 Å; Thermo Scientific) analytical columns (Acclaim PepMap TM RSLC: 75 μm x 75 cm, C18 2 μm, nanoViper) were used for the chromatographic separation at a 200 nl/min flow rate and 270 min gradient from 1 to 35% B (A= 0.1% FA in water, B= 0.1% FA in acetonitril). The mass spectrometer was operated in a data-dependent acquisition (DDA) mode using Orbitrap resolution in the MS1 (120k) and Ion Trap resolution in the MS2 with a 30% higher-energy collisional dissociation (HCD) for fragmentation.

MS/MS spectra were searched against the SwissProt (Mus musculus, release 2018_11) and contaminants database using MaxQuant v1.6.2.6 with the andromeda search engine (24). Searches were run against targeted and decoy databases. Search parameters included trypsin enzyme specificity, allowing for two missed cleavage sites, carbamidomethyl in cysteine as static modification and oxidation in methionine, acetyl in protein N-terminal and phosphorylation in Ser, Thr and Tyr as dynamic modifications. Peptide mass tolerance was 20 ppm, and the MS/MS tolerance was 0.6 Da. The minimal peptide length was 7 amino acids, the minimum score for modified peptides was 40, and the minimum delta score was 8. Peptide, protein and site identifications were filtered at a false discovery rate (FDR) of 1 % based on the number of hits against the reversed sequence database.

### Gene ontology

Protein expression enrichment analyses were computed using GSEA software (v4.1.0) (25). Samples were ranked according to the log2FC before submitted to GSEA. The enrichment analyses were run against the Ontology gene sets (c5) collection of the Molecular Signatures Database (MSigDB). An FDR value of 0.05 was used as cutoff for proteins positively regulated by p38α, and an FDR value of 0.25 for proteins negatively regulated by p38α, since no enriched GO terms passed the FDR<0.05 filtering criteria for these proteins in the KO/WT dataset, as per GSEA guidelines. Overlapped GO terms for KO/WT and PH/WT datasets were shown as bubble plots. DAVID database (v6.8) was applied to analyze gene Ontology enrichment in the phosphoproteomic dataset. Phosphoproteins containing phosphosites with a p-value< 0.05 and FC>1.5 were selected for GO analysis. A Benjamini-Hochberg correction value of 0.05 was used as cutoff. Overlapped GO terms in KO/WT and PH/WT were shown as bubble plots.

### Mapping functional scores and p38α substrates

The mouse phosphosites were firstly aligned to their conserved human phospho-sites using SITE_GRP_ID from PhosphositePlus (update: 2021/04/19). The functional score obtained (26) were assigned to the conserved phosphosites. Kinase_Substrates_Dataset from PhosphositePlus was used to map the p38α substrates. The overlapping was analyzed and visualized using the Python package.

### Identification of p38α interactors

To identify potential p38α protein interactors, we performed a bioinformatics analysis considering the information available in databases, the presence of docking (D-)motifs, and the modelling of docking peptides. In brief, we first retrieved all p38α interaction partners found in the IntAct database (27). We then used the experimentally determined D-motif of some p38α substrates (28–30) to derive four putative sequence motifs from a list of 48 known p38α substrates: (1) K..[RK].…L.L, (2) [KR].(4,5)[ILV].[ILV], (3) L..RR, and (4) [ILV](1,2).[RK](4,5). Then, we filtered out those interactors that did not have at least one putative D-motif. Overall, we retrieved 190 proteins as potential p38α interactors, 150 of which we classified as physical (i.e. identified in protein binding assays such as pull-downs, immunoprecipitations, and yeast two-hybrid screenings) and 50 as functional interactors (i.e. compiled from kinase and other enzymatic assays). The results were visualized using Cytoscape (v3.9.1).

### Kinase-Substrate Enrichment Analysis (KSEA)

The KSEA app (v1) was used to analyze the kinase activity. To extract known kinase-substrates relationships the database PhosphositePlus (PSP) was used. All conserved human phosphosites with log2 fold changes in our data were used to perform this analysis. The p-value represents the statistical assessment for the kinase activity (z-score). The chosen FDR cutoff (Benjamini–Hochberg correction) for significantly differential kinase activity was set at 0.05. Kinases with <5 substrates were excluded from the analysis.

#### Code and data availability

No custom codes were used. All software and code are available through the cited references. The MS proteomics data generated in this study have been deposited to the ProteomeXchange Consortium via the PRIDE partner repository (31) with the dataset identifier PXD034051 (Reviewer account details, Username; Password: FhD2I0hU). All relevant data are included in the article and/or supplementary information.

#### Statistical analysis

Data are expressed as mean ± SEM unless otherwise indicated in the figure legends. Statistical analysis was performed by using a two-tailed Student’s t-test with GraphPad Prism Software 8.02 (GraphPad Software, Inc., La Jolla CA). p-values are expressed as *p < 0.05, **p < 0.01, and ***p < 0.001.

## Results

### Identification of proteome and phosphoproteome changes upon p38α deletion or inhibition in cancer cells

We conducted quantitative proteomic and phosphoproteomic analyses to identify changes in protein expression and site-specific phosphorylations controlled by the p38 MAPK pathway in breast cancer cells. To this end, we used a cancer cell line derived from a mouse epithelial mammary tumor expressing the polyoma middle T (PyMT) antigen together with ubiquitin-CreERT2 (UbCre) and floxed alleles of *Mapk14*, the gene encoding p38α (19). This system allows the generation of a p38α KO upon 4-hydroxytamoxifen administration. To increase the confidence in the p38α-regulated proteomic and phosphoproteomic changes identified, we also targeted the p38α pathway using the inhibitor PH797804 (15). We confirmed decreased phosphorylation levels of the p38α pathway downstream components MK2 and Hsp27 in both p38α KO and PH797804-treated cells, as well as reduced p38α expression in p38α KO cells (Fig. 1A).

**Fig. 1.**
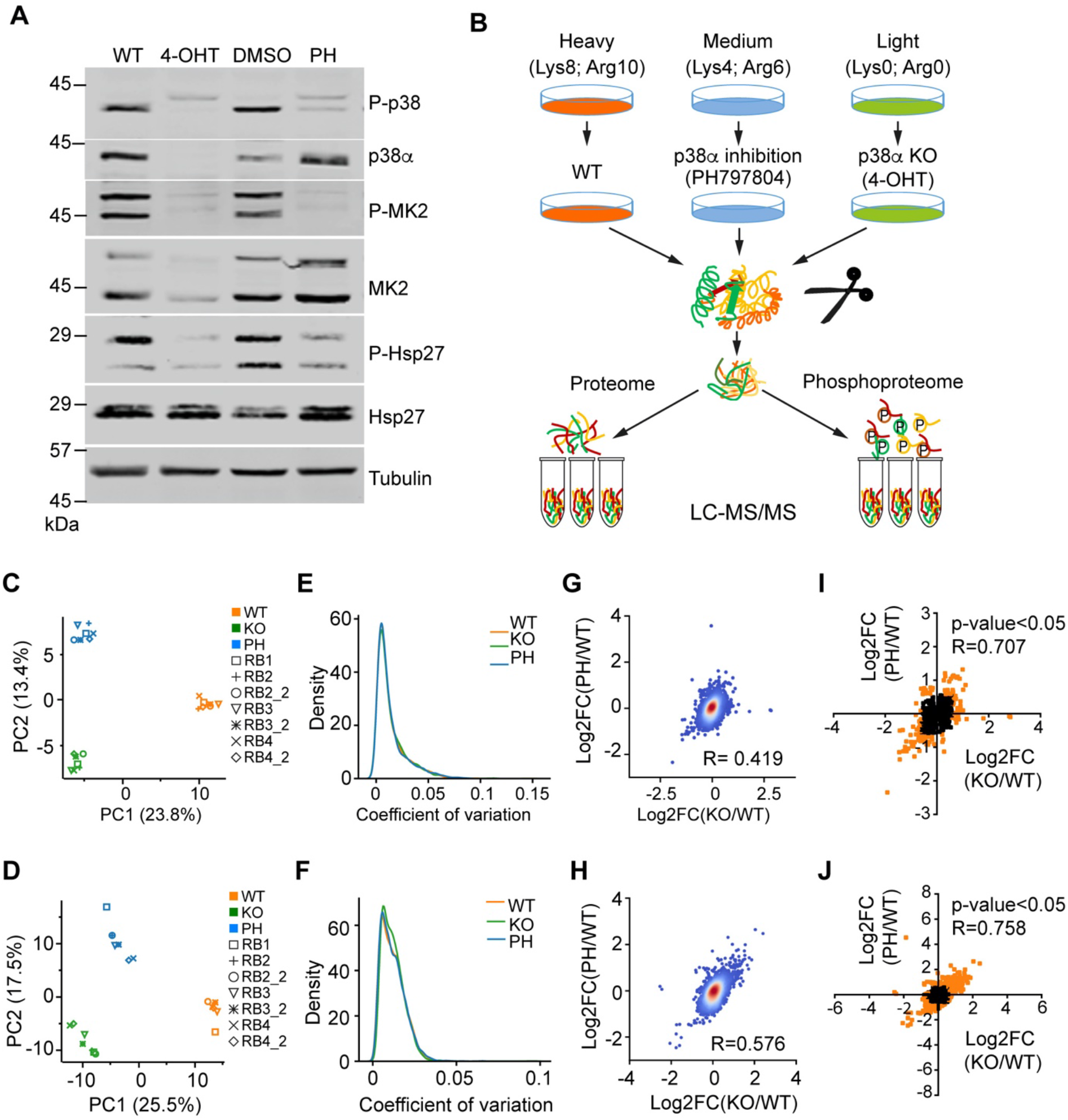
Preparation of samples and statistical analyses. *A*, cancer cells were incubated with 4-hydroxytamoxifen (4-OHT) for 48 h to induce the p38α KO and then left to recover for another 48 h, or were incubated with either the p38α inhibitor PH797804 (PH) or the vehicle DMSO for 24 h. Cell lysates were analyzed by immunoblotting using the indicated antibodies. *B*, schematic for the quantitative proteomic and phosphoproteomic analyses by SILAC. Cells were grown in culture media containing stable isotope heavy-, medium- and light-amino acids. Light-labeled cells were treated with 4-OHT, medium-labeled cells were treated with PH, and heavy-labeled cells were used as untreated controls (WT). Whole-cell lysates were digested and an aliquot was directly analyzed to identify the proteomic changes. The rest of the digested lysate was run through titanium oxide columns to enrich for phosphorylated peptides. Samples were analyzed by nano-liquid chromatography-electrospray ionization tandem mass spectrometry (nanoLC-ESI-MS/MS) in four biological replicates, of which three were re-analyzed as technical replicates. *C* and *D*, principal component analysis computed on protein intensities (*C*) and phosphosite intensities (*D). E* and *F*, the distribution of coefficient of variation for normalized intensity per protein (*E*) and per phosphosite (*F*) in the three samples. *G* and *H*, correlation between the log2 fold change (FC) of KO/WT and the PH/WT for all quantified proteins (*G*) and phosphosites (*H*). The color-coding indicates the density. Pearson coefficient correlation (R) is indicated. *I* and *J*, correlation between the log2FC of KO/WT and PH/WT of quantified proteins (*I*) and phosphosites (*J*) with *p*-value<0.05. Orange dots indicate FC>1.5. Pearson coefficient correlation is shown.

To quantify relative changes in the p38α-regulated proteome and phosphoproteome, we employed stable isotope labeling by amino acids in cell culture (SILAC) (32) (Fig. 1B), and calculated log2 fold changes and p-values of KO versus WT (KO/WT) and PH797804 versus WT (PH/WT) samples to facilitate subsequent analyses. We found that replicates of WT, KO and PH797804-treated samples clearly clustered together in a principal component analysis (Fig. 1C and D). Moreover, we confirmed a high quantitative reproducibility among biological replicates by the low coefficient of variation observed (Fig. 1E and F), validating the quality of the data. By plotting the correlations between KO/WT and PH/WT datasets, we estimated overall Pearson correlations of 0.419 and 0.576 for the proteomic and phosphoproteomic datasets, respectively (Fig. 1G and H). These values were further improved to 0.707 and 0.758, respectively, when we applied a p-value<0.05, suggesting a strong relationship between the changes observed upon p38α KO and inhibition (Fig. 1I and J). Taken together, these results indicate a good reproducibility of the quantitative analysis between the biological replicates, and that consistent changes are observed upon deletion or chemical inhibition of p38α.

The proteome analysis identified 3,920 proteins, of which 2,560 were successfully quantified, with 2,455 proteins being represented by at least two unique peptides (Fig. 2A and supplemental Table S1). The phosphoproteome analysis identified 6,726 phosphosites, of which 2,935 belonging to 1,202 proteins were quantified. About 92% of these phosphosites (2,690) were identified with a localization probability higher than 0.75, suggesting that the majority of the quantified phosphosites were localized with high confidence to specific positions on the phosphopeptide (Fig. 2B). We found that more than half of the quantified phosphoproteins (617) carried a single phosphosite, whereas 20%, 10% and 7% of the phosphoproteins contain two, three and four phosphosites, respectively (Fig. 2C and supplemental Table S2). We next examined the relative frequency of phosphoserine (pSer), phosphothreonine (pThr) and phosphotyrosine (pTyr) residues among the 2,935 quantified phosphosites. In agreement with previous studies (33), we mainly observed pSer (88%), followed by pThr (11%) and pTyr (1%) (Fig. 2D).

**Fig. 2.**
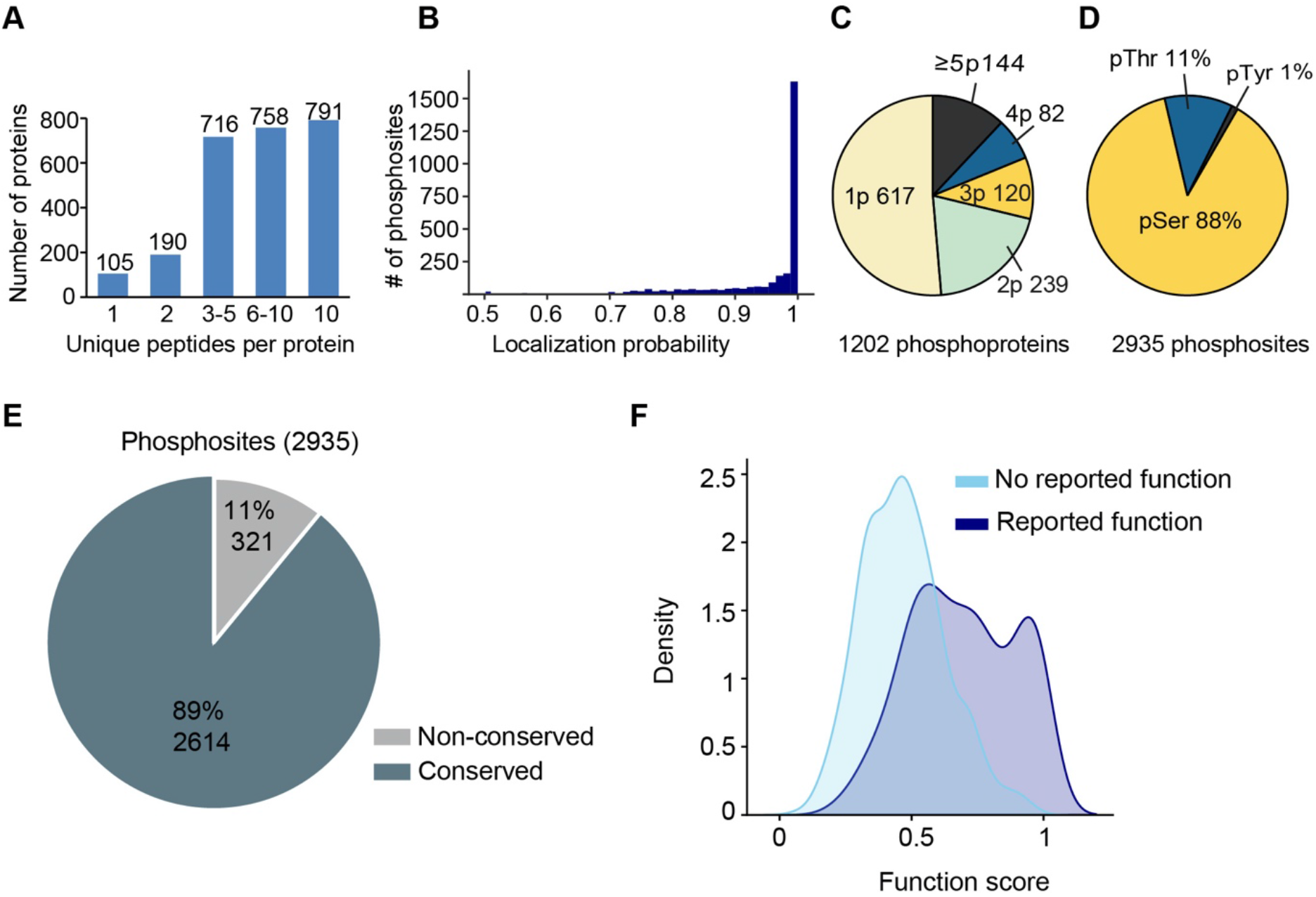
Characterization and functional importance of the phosphoproteome detected. *A*, bar chart showing the distribution of the identified proteins based on the number of unique peptides. *B*, histogram showing the distribution of the localization probability of the determined phosphosites in peptide sequences. *C*, pie chart showing the number of proteins containing the indicated number of phosphosites (p). *D*, pie chart showing the percentages of phosphorylated Serine(pSer), Threonine (pThr) and Tyrosine (pTyr) for all the quantified phosphosites. *E*, pie chart showing the results of mapping human orthologs for all the phosphosites quantified using the PhosphositePlus dataset. *F*, density plot showing the distribution of functional scores (26) for all the mapped phosphosites.

Since functionally important phosphorylation sites are more likely to be evolutionarily conserved, the mouse phosphosites were aligned to their conserved human orthologs using PhosphositePlus (5). We found that 89% of the quantified phosphosites (2,613) belonging to 1,131 proteins were conserved in human proteins (Fig. 2E). The conserved phosphosites were assigned functional scores (26) to identify those that were more likely to be relevant for cell fitness. The functional score distribution suggested that the conserved phosphosites tend to be functionally important, although many of them do not have reported functions yet (Fig. 2F).

### Proteins and phosphoproteins regulated by p38α in cancer cells

To identify the p38α regulated proteome and phosphoproteome, the datasets were filtered using a p-value < 0.05 and fold change (FC)> 1.5 (Fig. 3A and B). At the total protein level, we identified 43 proteins in p38α KO and 130 proteins in PH797804-treated cells that were downregulated compared with WT cells, with 25 proteins downregulated in both conditions. On the other hand, 37 proteins in p38α KO and 99 proteins in PH797804-treated cells were upregulated compared with WT cells, with 10 proteins upregulated in both cases (Fig. 3C and supplemental Table S3). Importantly, we confirmed the downregulation of the p38α protein in p38α KO cells but not in PH797804-treated cells, as expected from the immunoblotting results (see Fig. 1A). At the phosphorylation level, we identified 53 phosphosites on 45 proteins that were downregulated and 194 phosphosites on 134 proteins that were upregulated in p38α KO compared with WT cells. For PH797804-treated cells, 257 phosphosites on 146 proteins were downregulated and 155 phosphosites on 108 proteins were upregulated compared with WT cells. Similar to the protein expression data, there were about 5-fold more phosphosites downregulated in PH/WT than in KO/WT. Comparison of the two datasets identified 26 phosphosites on 21 proteins that were significantly downregulated and 88 phosphosites on 62 proteins that were significantly upregulated in both p38α KO and PH797804-treated cells versus WT cells (Fig. 3D and E, and supplemental Table S4), indicating that these phosphosites are very likely regulated by p38α.

**Fig. 3.**
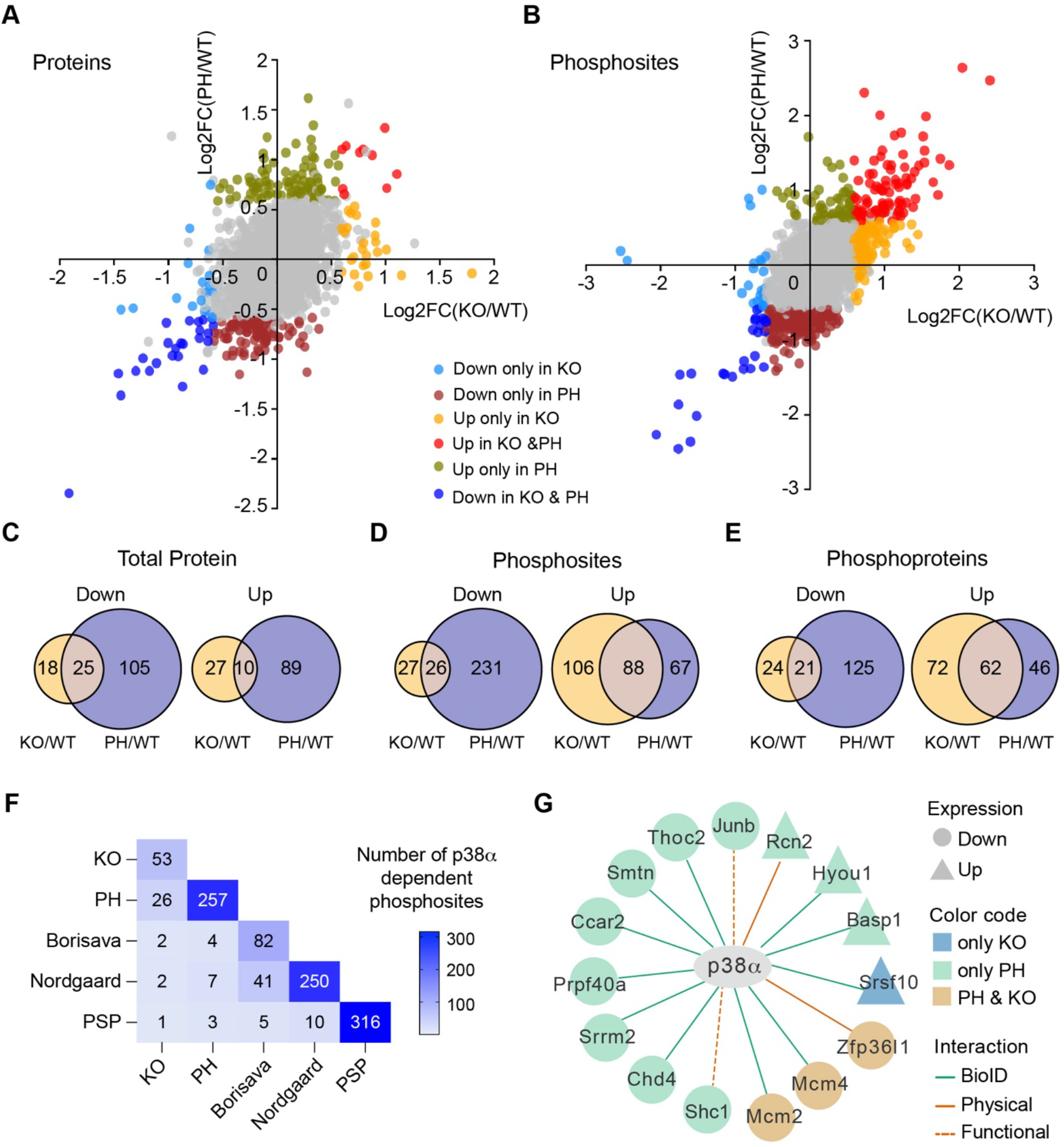
Quantification of the p38α-regulated proteome and phosphoproteome. *A* and *B*, scatter plots of log2 fold change (FC) of p38α KO/WT and the p38α inhibitor PH797804 (PH)/WT ratios for individual proteins (*A*) and phosphosites (*B*). Colors reflect protein or phosphosites that change significantly in both KO and PH (down: blue; up: pink), only in KO (down: skyblue; up: orange) or only in PH (down: dark red; up: olive), or that do not change in any of the samples (grey), using the thresholds of FC>1.5 and p-value<0.05. *C*-*E*, Venn diagrams showing the numbers of proteins (C), phosphosites (D) and phosphoproteins (*E*) that overlap between KO/WT and PH/WT (FC>1.5 and p-value<0.05) and that are either downregulated (Down) or upregulated (Up). *F*, overlap of p38α substrates listed in PhosphositePlus (PSP) and potential p38α targets previously reported by (14) or (13) with the p38α downregulated phosphosites identified in our KO/WT dataset (KO) or PH/WT dataset (PH). *G*, identification of p38α-regulated proteins that interact with p38α and could be potential substrates. The interactors were reported in the literature by using proximity-dependent biotinylation assays (BioID) or predicted according to protein binding assays (Physical) or kinase assays and other enzymatic studies (Functional). Shape indicates that the protein is significantly downregulated (circle) or upregulated (triangle) in KO/WT (blue), PH/WT (green) or both datasets (beig), using the thresholds of FC>1.5 and p-value<0.05.

As the phosphorylation of particular phosphosites is related to and can affect protein abundance, the overlap of the phosphoproteome and proteome datasets was determined. We detected total protein expression for around 80% of the downregulated phosphosites, and 58-66% of the upregulated phosphosites in either p38α KO or PH797804-treated cells (supplemental Fig. S1A). When we applied a cutoff with FC> 1.2 for protein expression changes, 15% of the phosphosites downregulated in p38α KO and 50% in the PH797804-treated cells were present in proteins that were also downregulated in those samples suggesting that differences in total protein levels could account for the changes in phosphorylation observed (supplemental Fig. S1B). However, only 2-7% of the downregulated phosphosites were present in proteins that were upregulated in the same samples. In addition, we found that over 25% of the upregulated phosphosites in either p38α KO or PH797804-treated cells corresponded to proteins that were downregulated (supplemental Fig. S1B), suggesting that they likely reflect real changes in phosphorylation, which might affect the abundance of those proteins.

Next, we compared the p38α-dependent phosphosites identified in p38α KO (53) or PH797804-treated cells (257) with previously known p38α substrates curated in the PhoshositePlus database (316) or reported in two phosphoproteomic studies (13,14) (82 in Borisova’s and 250 in Nordgaard’s) (Fig. 3F). Strikingly, out of the 26 phosphosites that we identified as very likely to be positively regulated by p38α (downregulated in both p38α KO and PH797804-treated cells) only 4 were previously reported: one (Mkl1_S492) in PhoshositePlus, and two each in Borisava’s (Hspb1_S15 and Rbm7_S136) and in Nordgaard’s (Hspb1_S15 and Nelfe_S115) datasets. For the phosphosites that were downregulated only in PH797804-treated cells, two were reported in PhoshositePlus (Ranbp2_S2505 and Trim28_S473), two in Borisava’s (Nelfe_S51, and Trim28_S473) and five in Nordgaard’s (Dnmt1_S140, Hspb1_S86, Nelfe_S51, Nop2_S59 and Trim28_S473) datasets. These results suggested that most of the p38α-regulated phosphosites identified in our analysis had not been reported before. This small overlap could reflect that previous studies mostly used stress stimuli, which likely shift the cellular response towards the phosphorylation of a limited number of substrates to respond to that particular insult, while in steady-state conditions p38α contributes to the fine-tuning of a larger number of processes (3). Consistent with this idea, the Borisova’s and Nordgaard’s datasets, which both focus on stress-regulated phosphorylations, show a higher overlap between themselves (Fig. 3F).

The interaction between protein kinases and their substrates is sometimes mediated by short linear motifs different from the phosphorylated peptide. As p38α is known to interact via docking motifs with the D domain of some of its substrates (28,34), we compared our dataset of 273 proteins that were significantly changed either in KO/WT or PH/WT (FC>1.5 and p-value<0.05), with a set of 381 potential p38α interactors identified using the BioID technique (35,36) or determined by *in-silico* analysis (see Experimental Procedures). We identified 15 potential p38α interactors in this group, which could therefore be potential p38α substrates (Fig. 3G). The majority of these interactors have not been reported to be phosphorylated by p38α, except for Shc1 (37). Zfp36l1 and Shc1 are also known targets of MK2 (38,39). It is worth mentioning that Basp1, Ccar2, Junb, Mcm2, Srrm2, Srsf10 were also identified in our phosphoproteomic analysis, and that Basp1, Mcm2, Junb and Srrm2 all changed in the same direction at both total protein and phosphorylation level upon p38α inhibition.

### Signaling networks regulated by p38α in cancer cells

A key determinant of kinase-substrate interaction is the recognition of phospho-acceptor residues and the surrounding linear sequence motif. Sequence analysis of the high confidence phosphopeptides regulated by p38α using iceLogo (40,41) showed that around 30% of the phosphosites (8 out of 26) that were consistently downregulated upon p38α KO or inhibition contained the Ser/Thr-Pro motif, which is present in many MAPK substrates and therefore could be directly phosphorylated by p38α. In addition, about 52% of these phosphosites (14 out of 26) showed a significant overrepresentation of Arg at the −3 position as well as the preference of hydrophobic amino acids at −5, −6, −7 positions, which resembled the optimal MK2 phosphorylation motif, Leu/Phe/Ile-x-x-x-Arg-Gln/Ser/Thr-Leu-pSer/pThr-Φ, with x denoting any amino acid and Φ a hydrophobic amino acid (42), suggesting an important contribution of MK2 (and perhaps the closely related kinase MK3) to the p38α regulated phosphoproteome (Fig. 4A and B). On the other hand, the 88 phosphosites that were negatively regulated by p38α showed a significant overrepresentation of Pro at +1 position and, to a lesser extent, Leu at −2, Thr at +2, Ala at +3 and Glu at +4, while Arg at −3 was clearly underrepresented (Fig. 4C and D). This points to the activation of proline-directed kinases upon p38α KO or inhibition, and the presence of Thr at +2 suggests the potential implication of other p38 MAPK family members, JNKs, ERK1/2 or CDKs as the kinases responsible for these phosphorylations.

**Fig. 4.**
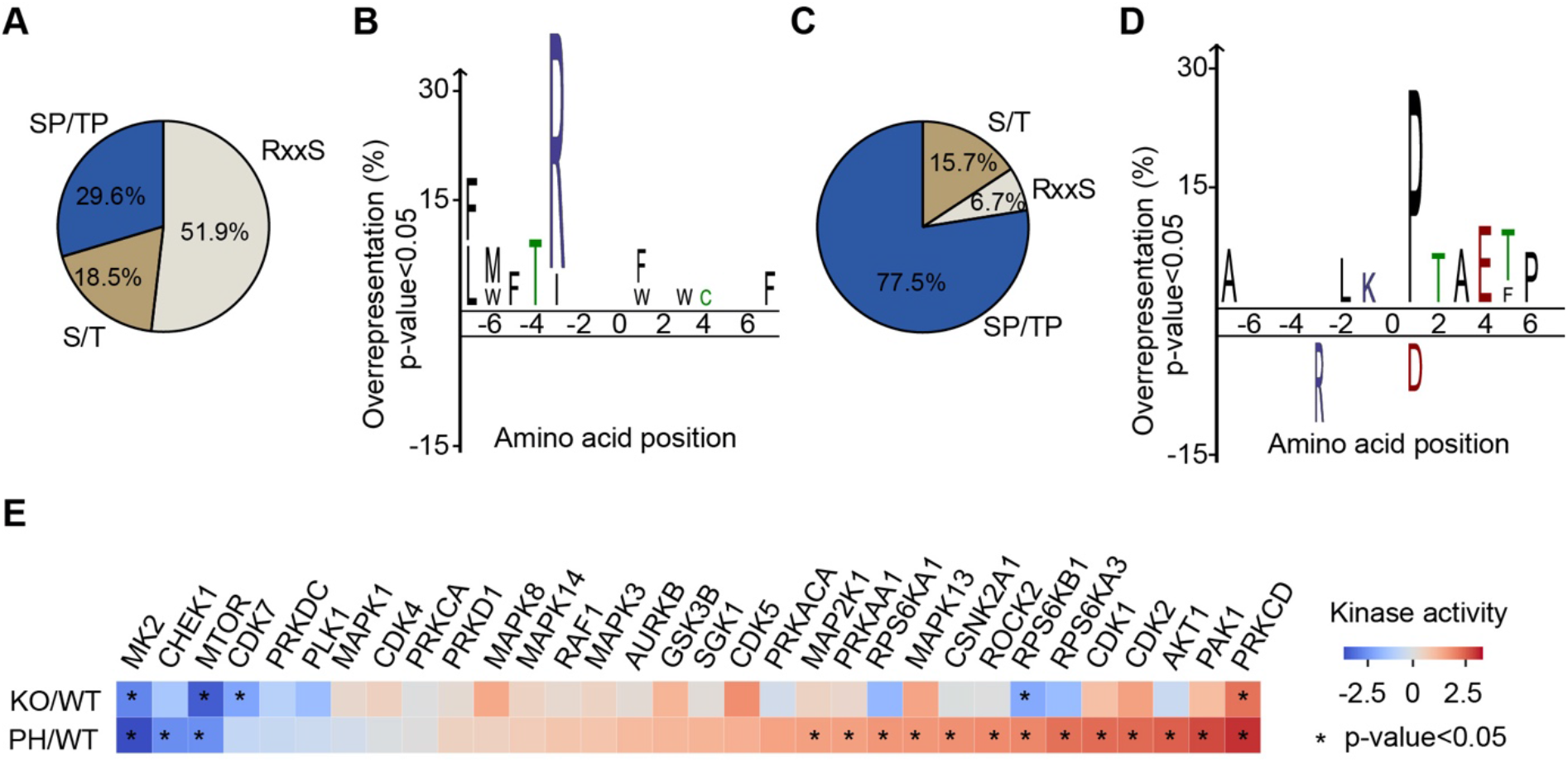
Characterization of the phosphoproteome modulated by p38α. *A* and *C*, pie charts showing the percentage of the indicated motifs in phosphosites positively (*A*) or negatively (*C*) regulated by p38α. *B* and *D*, sequence motif analysis using iceLogo of the phosphosites positively (*B*) or negatively (*D*) regulated by p38α. The frequencies of seven amino acids flanking the phosphorylated residue were compared with the frequencies in all the quantified phosphosites. Green, blue, red and black colors represent polar, basic, acidic and hydrophobic amino acids, respectively. *E*, kinase-substrate enrichment analysis (KSEA) using all the phosphosites that are conserved in human. Kinases with at least five potential substrates/ phosphosites are shown. Negative and positive values indicate that the corresponding kinase activity is predicted to be decreased and increased, respectively. Asterisk indicates p-value<0.05.

Sequence analysis indicated that over 50% of the phosphosites downregulated upon both p38α KO and inhibition were unlikely to be directly phosphorylated by p38α. Moreover, p38α KO or inhibition resulted in the upregulation of a substantial number of phosphopeptides, which again points to the implication of intermediary kinases. We used the Kinase-Substrate Enrichment Analysis (KSEA) to identify kinases that could phosphorylate the p38α-regulated phosphosites unlikely to be direct substrates (43–46). This analysis predicted that the activities of MK2 and mTOR were significantly decreased upon both p38α KO and inhibition (Fig. 4E). MK2 is a well-known downstream effector of p38α whose activity was expected to be reduced, validating the unbiased analysis performed (42,47). However, the identification of mTOR, a central regulator of mammalian metabolism (48), was more unexpected, since mTOR activity is not usually controlled by p38α, with a few exceptions (3,49). Our analysis also predicted decreased activities of the cell cycle regulatory kinases PLK1 and CDK7 in both p38α KO and PH797804-treated cells, although to a lesser extent. On the other hand, PRKCD was predicted to be significantly activated upon both p38α KO and inhibition (Fig. 4E).

Altogether, our results indicate that besides directly phosphorylating several proteins, p38α can also positively or negatively control a number of other kinases, which function as downstream effectors allowing signal spreading and integration with other signaling pathways in order to fine-tune specific cellular responses.

### p38α regulates DNA replication, RNA processing and cell adhesion in cancer cells

To identify processes regulated by p38α signaling, we performed functional enrichment analysis using the pre-ranked GSEA method (25,50). The enrichment results suggested that a large fraction of proteins positively regulated by p38α were involved in DNA replication and RNA metabolism, including mRNA translation and ribosome biogenesis (Fig. 5A). In contrast, the most enriched functions in the proteome negatively modulated by p38α were related to the extracellular matrix (ECM) and plasma membrane organization, which are linked to cell adhesion (Fig. 5B). Processes controlled by the p38α-regulated phosphoproteome were more challenging to predict given that proteins can be phosphorylated in several sites, which may differently modulate the protein activity. To address this, we analyzed the phosphoproteins regulated by p38α (FC>1.5 and p-value<0.05 in both KO/WT and PH/WT datasets) using the DAVID database (51). In agreement with the proteomics results, this analysis highlighted several processes related to cell adhesion (Fig. 5C). Since phosphorylation can modulate physical interactions between proteins and affect the complex stability and function, a protein-protein interaction (PPI) network (STRING score ≥0.4) was generated to illustrate physical or functional interactions among the p38α-regulated proteins and phosphoproteins that change significantly (FC>1.5, p-value<0.05). The biological function of the nodes was annotated in STRING. This network highlighted the implication of p38α in the regulation of cell adhesion and cytoskeleton, RNA processing and translation, and DNA replication (Fig. 5D).

**Fig. 5.**
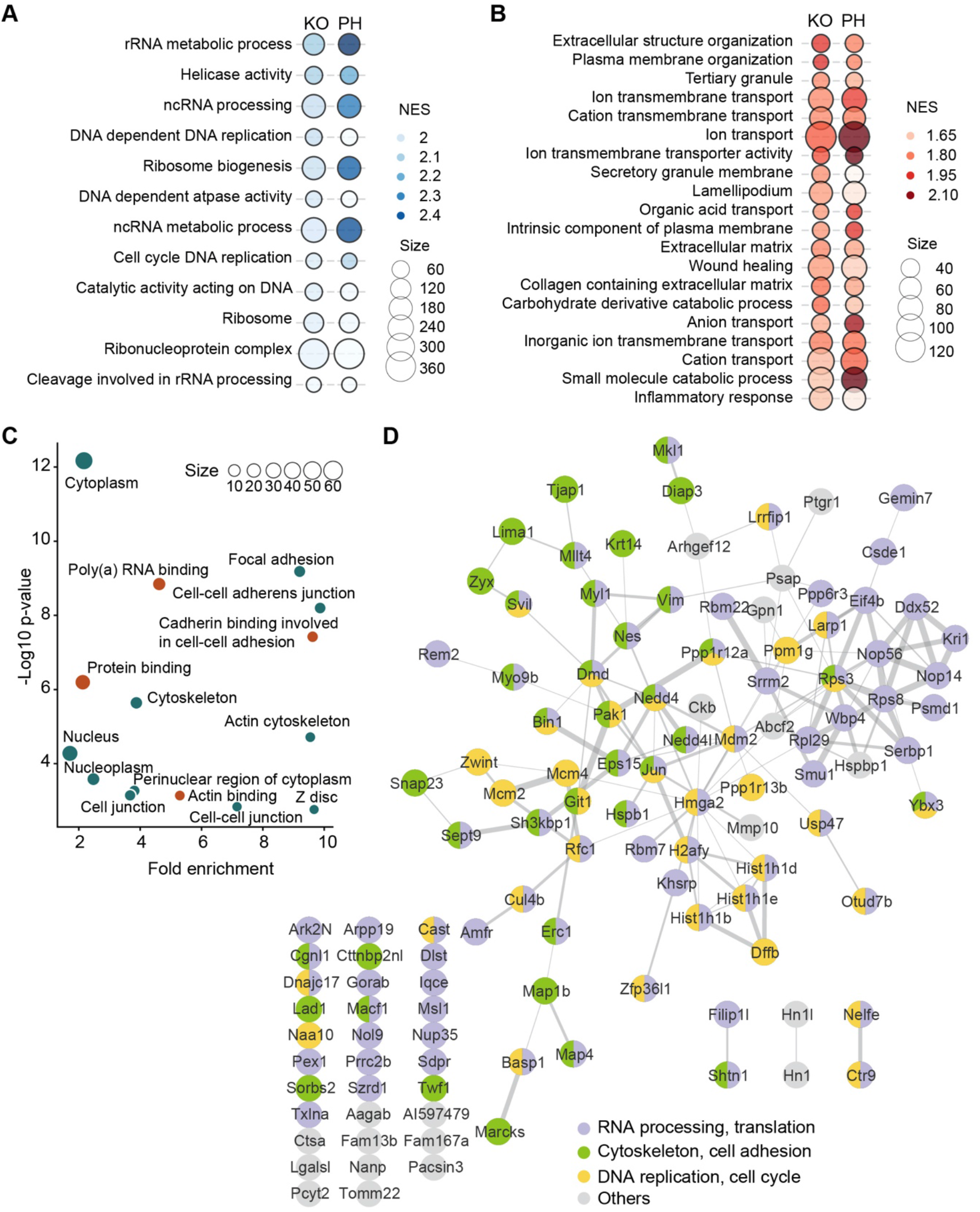
Functional analysis of the p38α-regulated proteome and phosphoproteome. *A*, biological processes represented in the proteins positively regulated by p38α. Gene ontology (GO) terms that passed FDR<0.05 and NES >1.9 thresholds are shown. *B*, biological processes represented in the proteins negatively regulated by p38α. GO terms that passed FDR<0.25 and NES >1.5 thresholds are shown. GO was analyzed using the pre-ranked GSEA method. FDR was estimated by the Benjamini-Hochberg procedure. NES stands for normalized enrichment score, and size indicates the number of genes involved in the corresponding biological process according to the input datasets. *C*, GO analysis for the phosphoproteins regulated by p38α using the DAVID database (FDR<0.05). Orange and green dots indicate molecular functions and cellular components, respectively. *D*, Functional signaling interaction networks modulated by proteins and phosphoproteins regulated by p38α. Networks were constructed using STRING (PPI scores≥0.4) and visualized in Cytoscape (v3.9.1).

Overall, the combined results of the proteomic and phosphoproteomic analyses identified the regulation of DNA replication, RNA processing and cell adhesion as important functions that are likely to be controlled by p38α signaling for cancer cell fitness.

### p38α negatively regulates cancer cell adhesion

Consistent with our predictions based on the omics analyses, p38α deletion in the breast cancer cell model used was previously shown to cause impaired DNA replication and the accumulation of DNA damage by affecting DNA repair signaling (19). Moreover, several studies have reported the implication of p38α and its downstream kinase MK2 in RNA processing, metabolism and stability (12,14,35,52,53).

However, the role of p38α in cell adhesion is poorly understood. Since many proteins and phosphoproteins in our dataset were related to cell adhesion (Fig. 5D), we decided to experimentally validate the implication of p38α in cancer cell adhesion. Using both noncoated and collagen-coated plates, which mimic cell adhesion to the ECM, we found that p38α KO or inhibition impaired cell adhesion (Fig. 6A). We also investigated cell detachment, as cell adhesion depends on the balance between attachment and detachment capabilities. After gentle trypsinization of cell monolayers, we found that fewer p38α KO cells remained attached on the plates compared to WT cells (Fig. 6B), further supporting that p38α facilitates cell adhesion.

**Fig. 6.**
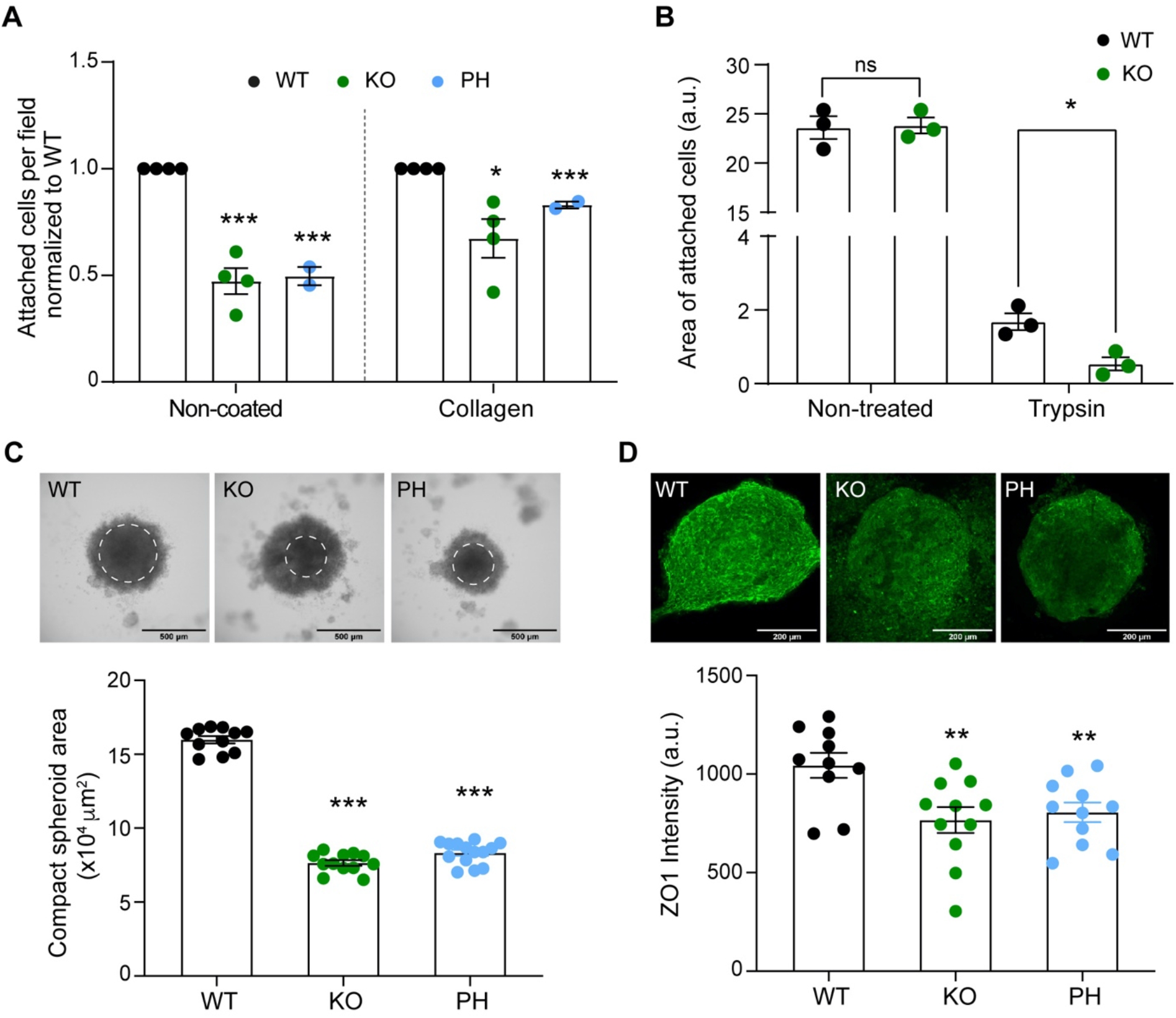
p38α negatively regulates cancer cell adhesion. *A*, cancer cells were incubated with 4-hydroxytamoxifen to induce the p38α KO, with the p38α inhibitor PH797804 (PH) or with the vehicle DMSO (WT) overnight, and then were cultured on either non-coated or collagen pre-coated plates. After 1 h, non-attached cells were removed and the cells that remained attached were fixed and stained for quantification. Numbers of attached cells were normalized to the WT plates in each condition. Attached WT cells vary between 10-20 in non-coated and 70-140 cells/field in collagen-coated plates. At least six different fields were analyzed in each case. n=2-4 biological replicates. *B*, cell monolayers of WT and p38α KO cells were treated with trypsin, washed and the whole area of cells remaining in the plates was quantified after crystal violet staining. n=3 biological replicates. *C*, spheroids were formed using the hanging drop method with WT or p38α KO cells, or with WT cells treated with PH for 48 h. The white circle indicates the compact area of the spheroids that was quantified. n=3 biological replicates. Data were obtained from 10-30 spheroids for each condition. Representative images are shown. *D*, spheroids prepared as in (*C*) were stained with anti-ZO-1 antibodies and the signal was quantified using the Fiji program. The quantification shows the ZO-1 mean intensity of z-stack images analyzed using z-Project sum slice function in Fiji. Representative confocal images are shown. n=3 biological replicates. Data were obtained from at least 10 spheroids for each condition. \**p* < 0.05, \*\**p* < 0.01, *** *p* < 0.001.

Besides adhering to the ECM, cells can also adhere to neighboring cells, using specialized structures, such as adherens junctions and tight junctions. Three-dimensional (3D) spheroids are considered useful *in vitro* models to study cell-cell adhesion in cancer cells as they may mimic more faithfully the *in vivo* situation in tumors than 2-D cultures (54). We found that WT cells consistently formed bigger and more compact spheroids than p38α KO cells or cells treated with PH797804 (Fig. 6C). Moreover, the impaired formation of spheroids correlated with decreased expression of the tight junction marker ZO-1 in p38α KO or PH797804-treated spheroids compared to the WT ones (Fig. 6D). Collectively, our results indicate that p38α can facilitate cell-cell adhesion in cancer cells.

### ArgBP2 mediates the regulation of cancer cell adhesion by p38α

Among the potential p38α targets regulating cell adhesion that were present in our Omics datasets, we focused on ArgBP2, an adaptor protein encoded by *Sorbs2* (Sorbin and SH3 domain-containing protein 2), which was consistently upregulated in p38α KO and PH797804-treated cells compared with WT cells. ArgBP2 has been reported to interact with several proteins involved in the regulation of actin (55) and to have a potential role in cell adhesion (56). We found that *Sorbs2* was upregulated upon p38α KO or inhibition, both at the mRNA and protein levels (Fig. 7A and B). Moreover, *Sorbs2* mRNA stability was similar between WT and p38α KO cells (Fig. 7C) but *Sorbs2* pre-mRNA levels were elevated upon p38α KO or inhibition (Fig. 7D). These results indicate that p38α can negatively control *Sorbs2* expression at the transcriptional level.

**Fig. 7.**
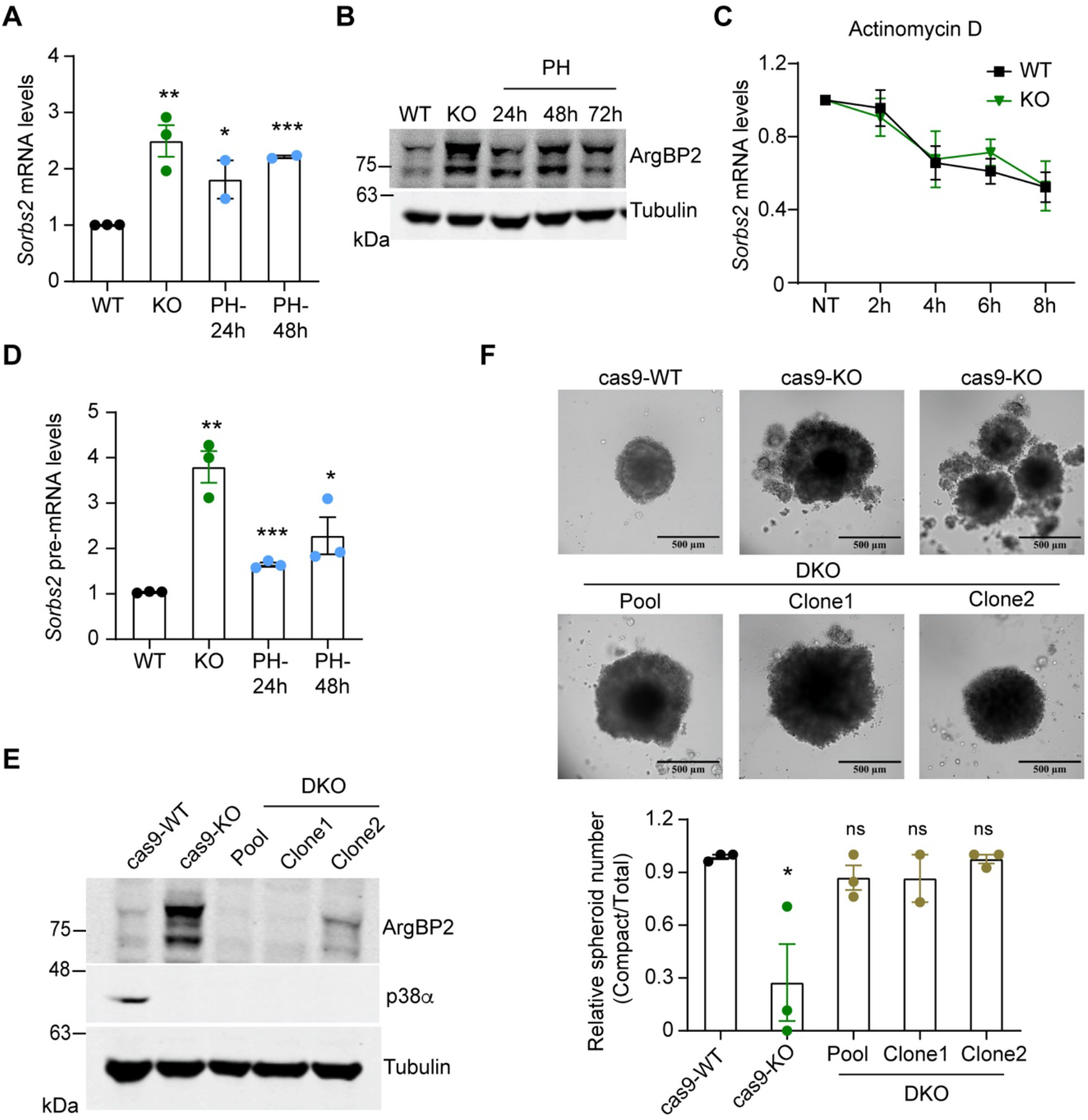
ArgBP2 is implicated in the regulation of cell-cell adhesion by p38α. *A*, the relative levels of *Sorbs2* mRNA encoding ArgBP2 were determined by qRT-PCR in cells WT, p38α KO or incubated with the p38α inhibitor PH797804 (PH) for the indicated times. *B*, the levels of the ArgBP2 protein were determined by immunoblotting in WT and p38α KO cells and in cells treated with PH for the indicated times. *C*, *Sorbs2* mRNA levels were determined by qRT-PCR in WT and p38α KO cells treated with the transcription inhibitor Actinomycin D (10 μg/ml), and were referred to the expression level in the untreated (NT) cells. Differences were not significant. *D*, the relative levels of *Sorbs2* pre-mRNA in WT and p38α KO cells were determined by qRT-PCR using primers that span an exon-intron junction. *E*, the expression of ArgBP2 and p38α in the indicated cell lines was determined by immunoblotting. Cas9-WT are cells expressing the Cas9 construct, which upon treatment with 4-hydroxytamoxifen (4-OHT) generate Cas9-KO cells. *F*, spheroids formed using the hanging drop method with cells WT, deficient in p38α (KO) or deficient in both p38α and ArgBP2 (DKO). The number of compact spheroids, defined by a diameter higher than 400 μm, was normalized to the total number of spheroids in each sample. ArgBP2 KO cells were obtained using the CRISPR/Cas9 system, and DKO cells were obtained by treating ArgBP2 KO cells with 4-OHT. Representative confocal images are shown, including two examples of p38α KO spheroids. n=3 biological replicates with at least 10 spheroids analyzed for each condition. Data are expressed as average ± SEM. \**p* < 0.05, ns, not significant.

To study the potential functional link between p38α and ArgBP2 in cancer cell adhesion, we downregulated both ArgBP2 and p38α and then performed spheroid formation assays (Fig. 7E). Strikingly, the less compact and sometimes fragmented spheroid phenotype observed with p38α KO cells was largely rescued by ArgBP2 downregulation, with the ArgBP2 and p38α double KO cells forming a high proportion of compact spheroids as observed with the WT cells (Fig. 7F). Taken together, our results support that enhanced ArgBP2 expression contributes to the impaired adhesion observed in p38α KO cells, indicating that p38α signaling controls cancer cell adhesion, at least partially, through the regulation of the adaptor protein ArgBP2.

## Discussion

We have used high-resolution quantitative mass spectrometry to study the p38α-regulated signaling networks in proliferating breast cancer cells. Previous reports using similar approaches have focused on the p38α targets engaged by particular stresses such as UV and anisomycin in U2OS cells (13,14). However, p38α can integrate a variety of intracellular signals in normally proliferating cells (3). Therefore, we decided to explore the targets and functions regulated by p38α, which could be potentially involved in cancer cell fitness, as these cells are endowed with rewired cellular signaling networks. Moreover, to improve the probability of identifying bona-fide p38α-regulated changes, we used complementary genetic and chemical approaches to downregulate p38α signaling, thus avoiding potential off-target effects due to the use of chemical inhibitors widely used in literature such as SB203580 (16–18), which can also inhibit both p38α and p38β with similar potency, making it challenging to distinguish p38α specific effects (1). These inhibitors have been also used to identify phosphoproteins potentially regulated by p38 MAPKs in differentiating myoblasts and in early mouse embryos (11,12).

Our studies identified with high confidence 114 phosphosites on 82 proteins that were regulated by p38α in steady state conditions. Most of them had not been previously linked to p38α, which is consistent with the idea that previous work was focused on the stress response, and the ability of p38α to orchestrate ad-hoc cellular responses relies on engaging a certain set of proteins required for a particular process to run. In contrast, proliferating cancer cells may use p38α in collaboration with other signaling pathways, to modulate the cell fitness, which probably involves a broader network of targets and processes. These may include other signaling proteins, such as additional kinases, so as to create a crosstalk that ultimately allows to maintain the fitness of the cancer cell. Consistent with this idea, we found that a large percentage of the p38α-regulated phosphosites actually increase upon p38α KO or inhibition, indicating that they are negatively regulated by p38α. Moreover, only 30% of the phosphosites that are positively regulated by p38α are predicted to be potential direct substrates. Collectively, these observations suggest a high degree of interplay between p38α signaling and other kinase pathways. A clear example of a kinase heavily involved in p38α signaling is MK2, whose potential substrates constitute the larger group of downregulated phosphosites upon p38α KO or inhibition. MK2 is known to form a complex with p38α in non-stimulated cells and contributes to its functions in various contexts including inflammation, DNA damage and survival to stress (42,57). Our results indicate that, in the absence of any stress stimuli, MK2 is also likely to mediate many p38α-driven responses.

Our analyses also predict the downregulation of Chk1 activity following p38α KO or inhibition, which fits well with previous studies showing the functional relationship between p38α and Chk1 in the DNA damage response to ensure genome stability during cell cycle progression (19,58). Moreover, we observed a number of changes that point to an important role of p38α in the regulation of RNA processing and ribosome biogenesis, two processes related to protein synthesis. These include the ribosomal proteins Rps3, Rps8 and Rpl29, the RNA helicase Ddx52, the nucleolar regulators of ribosomal assembly Nop14 and Nop56, the splicesome proteins Rbm22, Smu1, Smr2, Gemin7 or Wbp4, and the mRNA turnover regulators Khsrp, Larp1 and Zfp361. The implication of p38α in protein synthesis regulation has been ascribed in the literature mainly to the downstream kinases MK2, MNK1 and MNK2 (42,59), however, our results suggest and additional unanticipated layer of regulation that would involve the modulation of ribosome biogenesis. These observations are consistent with recent evidence showing that p38α can regulate ribosome-related gene expression, rRNA precursor processing, and polysome formation during blastocyst differentiation (12).

Altogether, our results provide new insights into the signaling networks regulated by p38α, and open the door to study the interplay with kinases whose activities we predicted to be regulated by p38α such as mTOR, Plk1 or Cdk7 both during cell homeostasis and in response to stress.

Besides the DNA damage response and RNA processing, which both have been previously linked to p38α in the literature, we identified cell adhesion as a function likely to be regulated by p38α signaling. Cell adhesion is key for cell viability as it is required not only for physically anchoring of the cells, but also for the integration of a variety of signals from the extracellular environment. We confirmed a role for p38α in cancer cell adhesion using several assays, most notably the formation of cell spheroids whose compactness is determined by homotypic cell-cell interactions. Our results are aligned with previous reports showing that p38α can control ZO-1 expression and the inhibition of p38α disrupts cell-cell junctions during blastocyst formation and in EMT (60,61), suggesting that p38α can control cell fitness through the modulation of intercellular junctions in homeostatic conditions. This probably becomes especially important in certain contexts where the proper establishment and maintenance of these junctions are likely to be critical such as during embryo development or tumor formation.

Mechanistically, we provide evidence that upregulation of ArgBP2 can mediate, at least partially, the effects of p38α on cancer cell adhesion. As an adaptor protein, ArgBP2 is important for cytoskeleton organization and cell adhesion (56), integrates the signals that control the balance between cell adhesion and motility (62), and has been suggested as a potential tumor suppressor in several cancer types (63–65). Interestingly, ArgBP2 expression has been inversely correlated with the expression of ZO-1 (66), which is consistent with our observation that ArgBP2 upregulation upon p38α inhibition correlates with decreased ZO-1 levels in spheroids. However, further work will be needed to know whether both observations are linked or they are independently controlled by p38α. Our omics analysis also identified several phosphoproteins regulated by p38α, such as Cttnbp2nl, Git1 and Zyz, which have been implicated in the process of cell adhesion through different mechanisms (67–69), as well as the kinase Pak1, a central regulator of the actin cytoskeleton, cell adhesion and motility (70). Interestingly, most of the phosphosites potentially related to cell adhesion identified in our analysis were not previously reported.

Taken together, our analyses reveal the complexity of the p38α-regulated signaling networks in proliferating cancer cells, and provide a valuable dataset of p38α-regulated proteomic and phosphoproteomic events in cancer cells to understand the versatility of this signaling pathway. Moreover, our work highlights the role of p38α in cancer cell adhesion and identifies ArgBP2 as a mediator of the p38α function in this process.

## Acknowledgments

We thank Isabel Fabregat, Judit López and Ester Bertran (IDIBELL, Barcelona) for discussions on experiments to analyze cell adhesion, Lidia Bardia and Anna Lladó of the IRB ADM core facility for help with confocal microscopy analysis, Eliandre de Oliveira and Antonia Odena of the PCB Proteomics platform for help with mass spectrometry sample preparation, and Joan Josep Bech for input early in the project. This work was supported by grants from the Spanish Ministerio de Ciencia e Innovación (MICINN, SAF2016-81043-R and PID2019-109521RB-I00) and AGAUR (2017 SRG-557) to A.R.N., and from Generalitat de Catalunya (RIS3CAT Emergents VEIS: 001-P-001647) and MICINN (PID2020-119535RB-I00) to P.A. Y. D. gratefully acknowledges financial support from the China Scholarship Council (NO. 201706310141), and N.R. and AF-T from Formación de Personal Investigador (FPI) predoctoral fellowships BES-2017-080105 and BES-2017-083053, respectively. The IRB Mass spectrometry and Proteomics core facility was supported by grant PRB3 (IPT17/0019-ISCIII-SGEFI/ERDF) and by the framework of the 2014-2020 ERDF Operational Programme in Catalonia (IU16-015983). IRB Barcelona is the recipient of institutional funding from MINECO through the Centres of Excellence Severo Ochoa award and from the CERCA Program of the Catalan Government.

---

**Supplemental Fig. S1.**
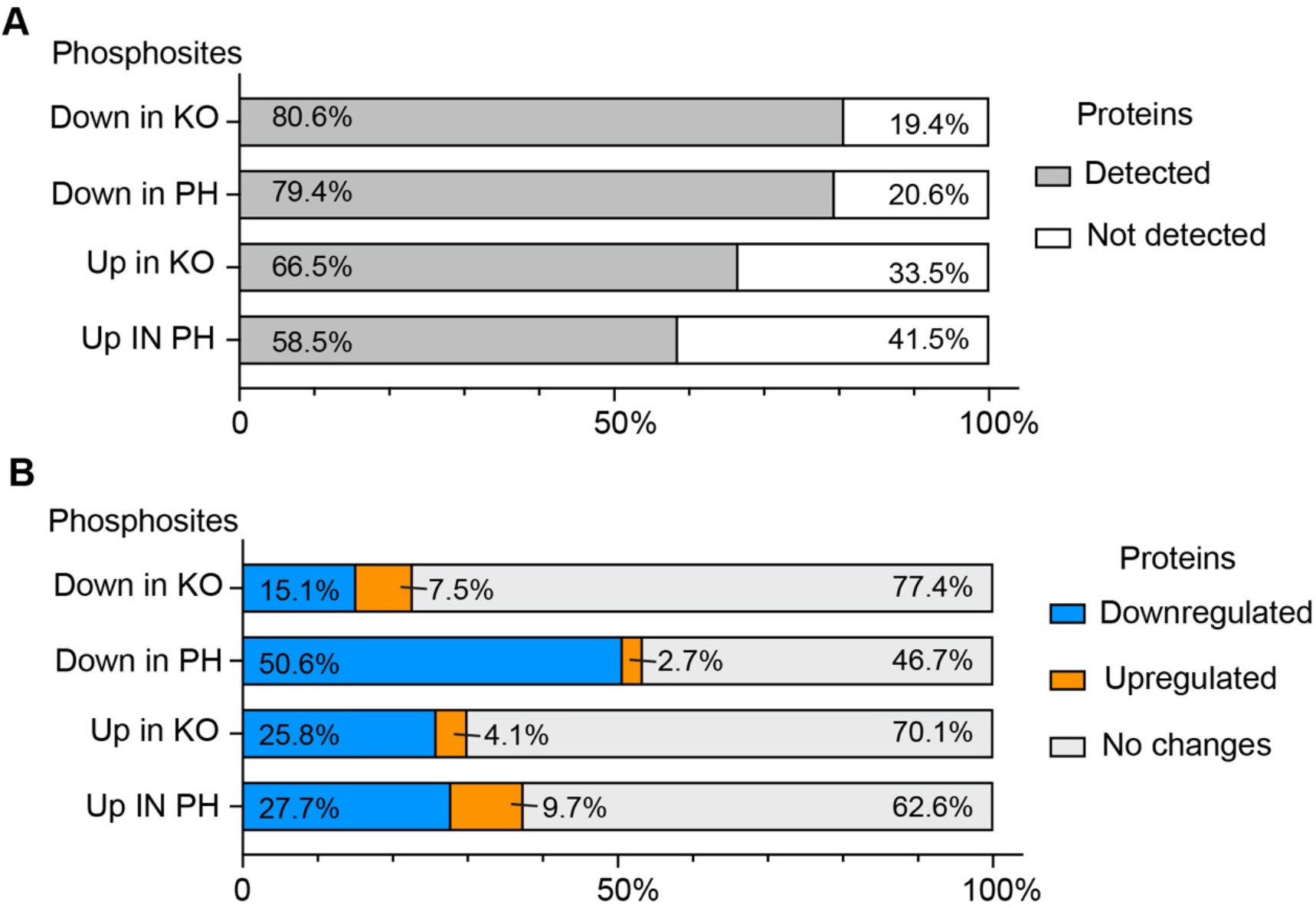
Overlap between the proteome and phosphoproteome datasets. *A*, percentage of phosphosites either Upregulated (Up) or downregulated (Down) in p38α KO or PH-797804 treated cells, and that are located in proteins detected or not detected in the proteome analysis. *B*, percentage of phosphosites that change in p38α KO or PH-797804 treated cell as indicated (Up or Down) and are located in proteins whose expression was either downregulated, upregulated or no changed. A cutoff of FC >1.2 in KO/WT or PH/WT was used to consider that the protein was downregulated or upregulated.

**Supplemental Table S1. Quantification of the total protein raw data** (*Excel Table*)

**Supplemental Table S2. Quantification of the total phosphosite raw data** (*Excel Table*)

**Supplemental Table S3. Proteins that change in both KO and PH-treated cells compared with WT cells** (*Excel Table*)

**Supplemental Table S4. Phosphosites that change in both KO and PH-treated cells compared with WT cells** (*Excel Table*)

